# Stable memory and computation in randomly rewiring neural networks

**DOI:** 10.1101/367011

**Authors:** Daniel Acker, Suzanne Paradis, Paul Miller

**Affiliations:** National Center for Behavioral Genomics, Brandeis University, 415 South St, Waltham, MA 02453; Volen National Center for Complex Systems, and University, 415 South St, Waltham, MA 02453; Department of Biology, Brandeis University, 415 South St, Waltham, MA 02453

## Abstract

Our brains must maintain a representation of the world over a period of time much longer than the typical lifetime of the biological components producing that representation. For example, recent research suggests that dendritic spines in the adult mouse hippocampus are transient with an average lifetime of approximately 10 days. If this is true, and if turnover is equally likely for all spines, approximately 95-percent of excitatory synapses onto a particular neuron will turn over within 30 days; however, a neuron’s receptive field can be relatively stable over this period. Here, we use computational modeling to ask how memories can persist in neural circuits such as the hippocampus and visual cortex in the face of synapse turnover. We demonstrate that Hebbian learning during replay of pre-synaptic activity patterns can integrate newly formed synapses into pre-existing memories. Further, we find that Hebbian learning during replay is sufficient to stabilize the receptive fields of hippocampal place cells in a model of the grid-cell-to-place-cell transformation in CA1 and of orientation-selective cells in a model of the center-surround-to-simple-cell transformation in V1. We also ask how synapse turnover affects memory in Hopfield networks with CA3-like, auto-associative properties. We find that attractors of Hopfield networks are remarkably stable if learning occurs during network reactivations. Together, these data suggest that a simple learning rule, correlative Hebbian plasticity of synaptic strengths, is sufficient to preserve neural representations in the face of synapse turnover, even in the absence of Hebbian structural plasticity.

## Introduction

Synapses are sites of communication between neurons. In mammals, dendritic spines are the major sites of excitatory synapses (Nimchinsky, Sabatini, & Svoboda, 2002). Dendritic spines in adult mammals can be eliminated or grow in new locations (Trachtenberg et al., 2002; Grutzendler, Kasthuri, & Gan, 2002). The reported degree of spine turnover in adult mice varies considerably by brain area. In cortex, most spines appear to be relatively stable once the critical period is over (Grutzendler, Kasthuri & Gan, 2002; Holtmaat et al., 2005; Attardo, Fitzgerald, & Schnitzer, 2015). In contrast, most spines in hippocampal area CA1 seem to be transient with a mean lifetime of approximately 10 days (Attardo, Fitzgerald, & Schnitzer, 2015; Pfeiffer et al., 2018). Thus, the hippocampus is expected to undergo dramatic structural remodeling with about 95% replacement of the CA1 spine population every 30 days (Attardo, Fitzgerald, & Schnitzer, 2015).

In light of such rapid and extensive restructuring of the hippocampal network, it is surprising that spatial memories can persist for at least one month in mice (Guskjolen, Josselyn, & Frankland, 2017). While these memories might not be entirely hippocampus dependent, Abraham and colleagues (2002) found that multi-afferent long-term potentiation (LTP) can persist for up to one year following stimulation in the rat dentate gyrus, suggesting that stable learning is possible within the hippocampus. Further, place fields of CA1 or dentate place cells can be stable for days to months in mice and rats (Thompson & Best, 1990; Agnihotri et al., 2004; Kentros et al., 2004; Ziv et al., 2013; Hainmueller & Bartos, 2018). While most place cells lose their location preference between exposures to an environment, some retain their place fields (Ziv et al., 2013; Rubin et al., 2015). The centroids of recurring place fields display little drift compared to their positions on earlier exposures (Ziv et al., 2013; Rubin et al., 2015). Most surprisingly, drift magnitude does not appear to increase over time, i.e. mean drift is no different after 30 days than after five days (Ziv et al., 2013). Similarly, the quality of a recalled spatial memory is not different between days one and 30 (Guskjolen, Josselyn, & Frankland, 2017).

Several proposed mechanisms might account for memory persistence in structurally unstable networks. One idea is that a subset of spines are stable, and these are sufficient to encode memory (Mongillo, Rumpel, & Loewenstein, 2016; Chambers & Rumpel, 2017). This hypothesis appears to hold true in some cortical regions, where a fraction of spines may persist throughout the lifetime of the animal (Yang, Pan, & Gan, 2009). A second possibility is that spine stability is regulated by activity-dependent structural plasticity (Bourjaily & Miller, 2011). Supporting this view, Hill and Zito (2013) found that LTP stabilizes nascent spines in hippocampal slices from neonatal mice. A third hypothesis is that network phenomena involving functional plasticity correct destabilization caused by random structural plasticity. One class of corrective functional plasticity involves feedback error signals (Chambers & Rumpel, 2017). Alternatively, Hebbian plasticity could lead to reinforcement of attractor states (Chambers & Rumpel, 2017), a process we investigate here.

Specifically, we use a modeling approach to explore the hypothesis that both memories and computations generating a neurons receptive field can be embedded at the network level in a manner that is robust to the incessant rearrangement of the connections between neurons (Mongillo, Rumpel, & Loewenstein, 2016; Chambers & Rumpel, 2017). We posit that the inevitable destabilization of circuits and resulting changes in neural responses caused by turnover of synapses is counteracted by Hebbian synaptic plasticity, particularly that which arises as a result of network dynamics during replay events.

The mechanism for stabilization relies on two general features essential to any type of pattern-stability and identity preservation in the midst of change. First, a degree of robustness is essential so that small changes do not destroy the pattern. In the case of neural receptive fields, the loss of a few inputs does not significantly alter a neuron’s response pattern because there is some redundancy among the original inputs. Second, naïve components must be able to respond to the state of the system and replace lost redundancy. Hebbian plasticity achieves this by specifically strengthening any new synapses from upstream neurons whose receptive fields overlap with that of the downstream neuron. Once a new synapse is strengthened, the corresponding upstream neuron supports the original receptive field by adding new redundancy so that the original receptive field can persist when more of the original synapses are lost.

To evaluate how effectively Hebbian plasticity of synaptic strengths could prevent drift in receptive fields due to connectivity changes, we measured the effects of synapse turnover on memory stability in a firing rate model of the grid-cell-to-place-cell transformation in CA1 (Brun et al., 2002, 2008; de Almeida, Idiart, & Lisman, 2009, 2012). Using this model, we demonstrate that learning during replay of grid cell activity stabilizes place fields over biologically relevant timescales in the presence of random synapse turnover. As grid cell replay may depend on the integrity of hippocampal replay originating in CA3 (Ólafsdóttir, Bush, & Barry, 2018), we also examined the effects of synapse turnover on the stability of stored attractors in CA3-like, sparse Hopfield networks whose within-circuit feedback connections lead to pattern completions (Hopfield, 1982; Wittenburg, Sullivan, & Tsien, 2002; de Almeida, Idiart, & Lisman, 2007). We show that learning during network reactivations allows stable activity patterns of the feedback circuit to survive complete rearrangement of the connection matrix. Together, our results suggest that long-term, hippocampal memories may persist despite on-going synapse turnover.

Finally, we asked whether our model can account for functional stability in non-hippocampal neural networks undergoing synapse turnover. In primary visual cortex (V1), orientation preference maps appear to be stable over time (Stevens et al., 2013), while dendritic spines (Holtmaat, Trachtenberg, & Wilbrecht, 2005; Tropea et al., 2010; Yu, Majewska, & Sur 2011) and axonal boutons (Stettler et al., 2006) turn over throughout the life of the animal. To evaluate the effect of activity-independent rewiring of LGN-to-V1 synapses on orientation tuning stability, we simulated the center-surround-cell-to-simple-cell transformation in V1. In this model, we observed that orientation preferences in individual V1 cells survive complete rearrangement of input connections when visual stimuli were paired with Hebbian learning. Taken together, our results suggest mechanisms by which activity patterns can remain stable, thus maintaining a stability of identity at the single-neuron level, even when all of the connections providing input to the neuron have changed.

## Materials and Methods

We use four separate neural firing rate models to test our hypothesis. Model one and model two represent the transformation of grid cell inputs from entorhinal cortex (through the temporoammonic tract and perforant path) into the place fields of CA1 pyramidal cells. Model one includes only a single pyramidal cell in CA1, while model two expands the representation of CA1 to include 2000 pyramidal cells and feed-back inhibition. Model three is an associative memory network representing internal excitatory and inhibitory connections in CA3. Model four represents the transformation of center-surround cell inputs from lateral geniculate nucleus (LGN) into the orientation tuning of V1 simple cells.

### Models one and two—grid cell to place cell transformation

#### i. Grid cells

Data were simulated by assuming a one-meter linear enclosure divided into one-centimeter bins. The activity of each cell was characterized by its firing rate in each bin. We simulated a library of 10000 grid cell responses according to a method described by Blair et al. (2007) (Eq. 1). Using this method, a grid cell’s firing rate varies in a hexagonal grid across the enclosure, and the orientation, phase, and offset of the grid are tunable parameters.

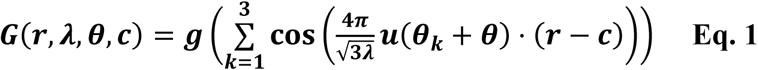

In equation one, *G* is a grid cell's firing rate, *r* is the animal’s position in two-dimensional space, *λ* is the distance between grid vertices and ranged from 30 to 100 centimeters, *θ* is the angular offset and ranged from 0° to 60°, and *c* is the offset in two-dimensional space and ranged from zero to 100 centimeters in both dimensions. *g* is a gain function *g*(*x*) = exp[*a*(*x* − *b*)] − 1, where *a* modulates the spatial decay and was set to 0.3, and *b* modulates the minimum firing rate and was set to −3/2. The hexagonal grid is created by summing cosine gratings angled at *θ*_1_ = −30°, *θ*_2_ = 30°, and *θ*_3_ = 90°. *u* is the function *u*(*θ_k_*) = (cos(*θ_k_*), sin(*θ_k_*)). Grid parameters were assumed to be the same as for grid cells at the same position on the dorsal-ventral axis innervating dentate (Moser, Moser, & Roudi, 2014) and were adapted from de Almeida, Idiart, & Lisman (2009).

#### ii. Place cells

We simulated place cells as described by de Almeida, Idiart, & Lisman (2009). Each cell received and summed excitatory input from 1200 randomly selected grid cells out of the library of 10000 grid cells. This number of grid cell inputs was chosen to approximate the total number of spatially modulated cells from entorhinal cortex that form synapses on the spines of a CA1 pyramidal cell. CA1 pyramidal neurons receive synaptic input from layer III of entorhinal cortex on dendrites in stratum lacunosum-moleculare. The number of inputs, 1200, was obtained by taking the average number of spines per CA1 pyramidal cell on dendrites in stratum lacunosum-moleculare (approximately 1500 spines [Megías et al., 2001]) and multiplying by the fraction of neurons with spatial tuning in layer III of entorhinal cortex (about 80%, [Sargolini et al., 2006]). We disregarded non-cortical inputs because place fields can be observed in CA1 after the removal of all input from CA3 (Brun et al., 2002).

The activity of each place cell was determined in each one-centimeter bin of the one-meter linear enclosure. The sum of synaptic input to a place cell was calculated as

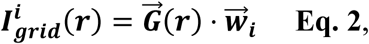

where 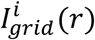 is the input to the *i^th^* place cell at position *r*, 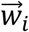 is the synaptic strength vector representing grid cell synapses onto the *i^th^* place cell, and 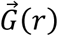 is the vector of all grid cell firing rates at position *r*.

In the case of model one where there was only one place cell, the firing rate was given by 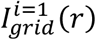, which is the sum of synaptic inputs to the place cell at position *r* as shown in equation three. Further, the place cell’s firing rate was set to zero at all locations except for the ten where excitation was greatest. This procedure was not intended to be realistic, but to assure that the single place cell in model one consistently had a defined place field. A more realistic approach to the same problem is used in model two.

In model two, there were 2000 place cells. In this case, place cell firing rates were modulated by feed-back inhibition. We assume that feed-back inhibition is triggered by a single place cell spike and is strong enough to suppress place cell firing (de Almeida, Idiart, & Lisman 2009). Under these assumptions, a place cell cannot fire unless it spikes before receiving feed-back inhibition (de Almeida, Idiart, & Lisman 2009). This process can be approximated by a rule stating that a cell can only fire if it is excited to within some fraction (***k***) of the excitation received by the most excited cell (de Almeida, Idiart, & Lisman 2009). Therefore, the firing rate of a place cell was calculated as

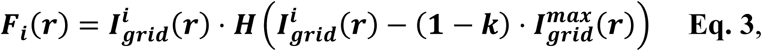

where *F_i_*(*r*) is the place cell’s firing rate at position *r, H* is the Heaviside function, and 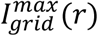 is the sum of excitatory input received by the most strongly excited place cell at position *r*. The parameter *k* is the fraction of 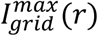 to which a place cell must be excited in order to fire. We set *k* = 0.10, the approximate ratio of feed-back delay (the time between a pyramidal cell spike and the arrival of feed-back inhibition [2 to 3 milliseconds] [Miles, 1990]) to the membrane time constant (about 23 milliseconds in pyramidal cells [Turner & Schwartzkroin, 1983]) (de Almeida, Idiart, & Lisman 2009, 2012).

#### iii. Synaptic strengths and learning

Synaptic strength was assumed to be a function of synaptic size (Eq. 4 [de Almeida, Idiart, & Lisman 2009]), and the distribution of synaptic sizes was assumed to be the same as that empirically determined for dentate (Eq. 5) (de Almeida, Idiart, & Lisman 2009, 2012).

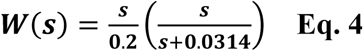

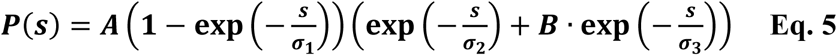

In equations four and five, *W*(*s*) is the synaptic strength of a synapse of size *s*. The synaptic size was an area value that could range from zero to 0.2 square microns. *P(s)* is the probability density function of the distribution of synaptic sizes. The values of constants were obtained from Trommald & Hulleberg (1997) and were *A =* 100.7, *B* = 0.02, *σ*_1_ = 0.022 square microns, *σ*_2_ = 0.018 square microns, and *σ*_3_ = 0.15 square microns.

Synaptic strengths were updated once per running session within the linear enclosure. The strengths of realized synapses were updated according to Hebb’s rule (Eq. 6). After each strength update, we scaled the total strength of synapses converging onto each place cell by dividing by the sum of synaptic strengths and multiplying by the expected value of the sum of 1200 random draws from the empirical distribution of synaptic strengths. This procedure represents the homeostatic scaling response seen in biological neurons (Turrigiano, 2010).

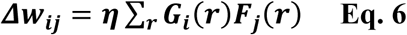

In equation six, *w_ij_* is the strength of the synapse connecting grid cell *i* and place cell *j*, *G_i_*(*r*) is the firing rate of grid cell *i* at position *r*, and *F_j_*(*r*) is the firing rate of place cell *j* at position *r*. The learning rate *η* influences the relative contribution of the pre-existing synaptic strength and the change in strength induced by learning. At high values of *η*, the change in strength will eclipse the preexisting strength, although scaling will cause the overall distribution of strengths to be largely unchanged. Unless otherwise stated, the learning rate was 1e-4, as determined in figure 3A.

#### iv. Synapse turnover

Synapse turnover occurred prior to each running session in the linear enclosure. In the case of model one, between 10% and 100% of the active grid cell inputs were replaced. In the case of model two, a random 114 of the 1200 active grid cell inputs per place cell were replaced. The number 114 corresponds to *N*_0_ – *N*(*T*) in the exponential decay model (Eq. 7) assuming a mean synapse lifetime (*τ*) of 10 days (Attardo, Fitzgerald, & Schnitzer, 2015) with the inter-trial time (*T*) taken to be one day. This is derived from the exponential decay model

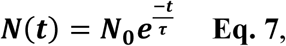

where *N*(*t*) is the number of original synapses remaining after *t* days, *N*_0_ = 1200 is the number of synapses on day zero, and *t* = *T*. Synapse turnover resulted in the erasure of learned synaptic strengths, with newly formed synapses taking random values sampled from the empirically determined distribution described above.

#### v. Place field analysis

Place cells were said to have a place field on a given trial if there was exactly one continuous region in the enclosure, at least five centimeters in length, where their firing rate was within 80% of their maximum firing rate on the same trial. The place field drift on a given trial (Ω*_t_*) was calculated for all place cells with place fields both on trial *t* and trial zero as the absolute centroid offset Ω*_t_* = |*C_t_* − *C*_0_|, where *C_t_* is the cell’s place field centroid position on trial *t* and *C*_0_ is the position on day zero.

### Model three—maintenance of recurrent attractor states

We modeled sparsely connected, asymmetric attractor networks based on the classic Hopfield model (Hopfield, 1982). Briefly, networks consisted of internally connected units with a globally defined probability of connection between any two units. Self-loops were allowed. Synaptic strengths could take on both positive and negative values, with negative values representing feed-forward inhibition through intervening inhibitory interneurons (Wittenburg, Sullivan, & Tsien, 2002).

#### i. Dynamics

Networks were simulated according to Wittenburg, Sullivan, & Tsien’s (2002) equations 1 and 2, modified for biological realism to permit only positive firing rates. The modified equations are shown here as equations eight and nine.

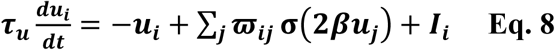

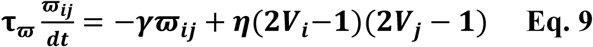

Equation eight describes how neuron *i*’s membrane potential (*u_i_*) changes across iterations of the network during a single activation. The term − *u_i_* causes the membrane potential to decay to zero in the absence of input. The term Σ*_j_ ϖ_ij_ σ*(2*βu_j_*) describes the change in membrane potential due to synaptic input, where *σ* is the logistic function *σ*(*x*) = 1/[1 + exp(−*x*)]. We set *β* = 1. The term *I_i_* represents external input and was set to zero except during training. In cases where one pattern was stored in the network, the membrane time constant (*τ_u_*), the synaptic strength time constant (*τ_ϖ_*), the learning rate (*η*), and the synaptic strength decay constant (*γ*) were set to one. In cases where multiple patterns were stored, the synaptic strength time constant (*τ_ϖ_*) was set to 1000 during training and to 100 during reactivations.

Equation nine describes how the synaptic strengths change between activations of the network. During reactivations, the change in strength for the synapse from the *j^th^* to the *i^th^* neuron (Δ*ϖ_ij_*) was calculated on each activation after 12 time steps. During training of multiple patterns, Δ*ϖ_ij_* was calculated on every activation. The term −γ*ϖ_ij_* causes the synaptic strengths to decay toward zero. This simulates the decay of synaptic receptors (Wittenburg, Sullivan, & Tsien, 2002). The term *η*(2*V_i_* − 1)(2*V_j_*·− 1) describes the Hebbian change in synaptic strength at realized synapses, where *η* is the learning rate, and *V_i_* = *σ*(2*βu_i_*) is the firing rate of neuron *i*.

#### ii. Training

In cases where one pattern was stored, training was performed by setting the synaptic strength matrix of internal connections to *ϖ* = (2*Y_goal_* − 1)*^T^* (2*Y_goal_* − 1) × *C*, where *Y_goal_* is the binary, {0, 1}, matrix of target firing rates (rows of *Y_goal_* represent individual, unraveled patterns; columns of *Y_goal_* represent the firing rates of individual cells), *C* is matrix of internal connections with values in (0, 1), and × represents elementwise multiplication.

In cases where multiple patterns were stored, the network was initialized with the neutral synaptic strengths (all strengths were set to zero) and membrane potentials were initially set to zero. A strong external input (*I_i_* in equation seven) in [−80, 80] was provided to each unit to drive the network towards a target attractor state. This ensured that external input dominated network activity during training. The network was updated according to equation eight for a number of time steps equal to 1200 times the number of patterns being trained. This is similar to the procedure used by Wittenburg, Sullivan, & Tsien (2002) to store multiple patterns in fully connected Hopfield networks. The pattern setting the external input variable was changed every 12 time steps to the next pattern in the full sequence being trained. After every time step, membrane potentials were supplied to equation nine to update synaptic strengths.

#### iii. Reactivation

In cases where one pattern was stored, reactivations were performed by initializing the network with random membrane potentials in the range (−0.004, 0.004) and no external input. In cases where multiple patterns were stored reactivations were performed as described above except that membrane potentials were initialized so as to put the network in the vicinity of the attractor for a specific stored pattern. Initial membrane potential values for these reactivations were obtained by taking a pattern used for training, adding uniform random noise in the range (−2.5, 2.5) to each value, and taking the hyperbolic tangent of the pattern plus noise. This procedure was performed on each reactivation so that the network initial states varied across reactivations of the same pattern. The Pearson’s correlations of these initial states with the training patterns, on average, were 0.5 for the corresponding pattern and zero for unrelated patterns. After each reactivation, the pattern was changed to the next in the full sequence of patterns. The full sequence of patterns was repeated for a variable number of cycles.

In each case, the network was updated for 12 iterations after each initialization according to equation seven. Membrane potentials on the 12^th^ iteration were supplied to equation nine to update synaptic strengths before the next activation. Synapse turnover occurred at the beginning of each reactivation. During turnover, a specified percentage of synapses was replaced. Synapse turnover resulted in the erasure of learned synaptic strengths, with newly formed synapses taking on random values in the range (−1, 1).

#### iv. Attractor integrity analysis

The quality of an attractor for a specific pattern at a given activation of the network was assessed as the Pearson’s correlation of the vector of firing rates on the final iteration of that activation with the target normalized firing rate vector (the first row of *Y_goal_*) used for training that pattern. To assess the change in memory quality over time, we compared the attractor integrity values of individual networks on reactivations one and 100. To derive the confidence intervals shown in Figure 10, we performed locally weighted regression using the “loess” function in the R “stats” package (R Core Team, 2017) with span set to 0.25 and otherwise default parameters. To estimate integrity scores at missing combinations of turnover rate and network in-degree, as shown in Figure 11, we performed bilinear interpolation using the “interp” function in the R “akima” package (Akima & Gebhardt, 2016; R Core Team, 2017) with the square-root of in-degree as the *x* parameter, turnover rate as the *y* parameter, a 40 × 50-pixel grid size, extrapolation outside of the convex hull of input data, averaging of duplicate parameter combinations, and otherwise default parameters.

### Model four—LGN inputs to orientation-selective neurons in V1

#### i. Grating stimulus

Data were simulated by assuming a square visual field. Visual input consisted of a series of static gratings presented at a variety of orientations and phases. The grating was defined with 100 × 100-pixel resolution. The luminosity at each pixel was determined as

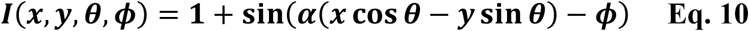

where *x* and *y* are a grid of values covering the range (−1, 1) in both dimensions. The grid orientation, *θ*, was sampled evenly in the range of zero to 180-degrees, and the grid offset, *ϕ*, was sampled evenly in the range of zero to *2π* − 0.1 in steps of 0.1. The parameter *α* controls the spatial period and was set to 10.

#### ii. LGN center-surround cells

We simulated the activities of 20000 LGN center-surround cells (10000 on-center and 10000 off-center). The input to each LGN cell was calculated at each position according to a difference-of-Gaussians function (Eq. 11).

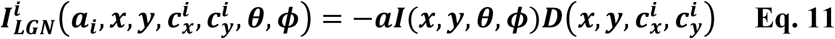

In equation 11, 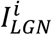 is the input to LGN cell *i* at position *x, y* (pixel units). The parameter *a_i_* is positive-one for on-center cells and negative-one for off-center cells. The parameters 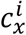, and 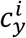 are the x and y coordinates of the receptive field centroid of cell *i*. Receptive field centroids were randomly chosen positions in the visual field. G is the Gaussian 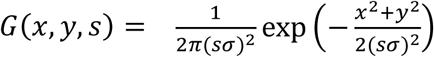, where *σ* is the standard deviation, set to 5 pixels, and the parameter *s* scales the standard deviation. D is the difference-of-Gaussians 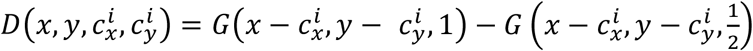, where the standard deviation of the inner Gaussian is half that of the outer.

The firing rate of each LGN cell was determined at each orientation and phase of the grating stimulus as the rectified sum of input across all positions on the visual field (Eq. 12).

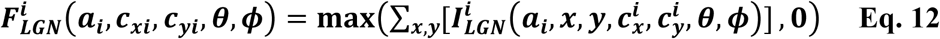

#### iii. V1 simple cells

We simulated the activities of 1000 V1 simple cells. Real V1 cells are estimated to receive between 50 and 150 inputs from LGN cells (Van Hooser et al., 2014). In our model, V1 cells initially received synaptic input from 130 LGN cells. Simple cells in tree shrews are characterized by a single dark receptive field flanked by two light fields (Lee, Huang, & Fitzpatrick, 2016). Therefore, we initial arranged on- and off-center LGN inputs to minimize the mean Euclidean distance between LGN cell receptive field centroids and an “off” bar flanked by two parallel “on” bars across the visual field. For each V1 cell, bars were spaced 15 pixels apart. The “off” bar was 60 pixels in length, and the “on” bars were 40 pixels in length. Bars were centered on the randomly positioned receptive field centroid of the V1 cell. Bars were oriented at a randomly chosen angle of between zero and 180-degrees. Thirty-three on-center LGN inputs were aligned to each “on” bar, and 64 off-center LGN inputs were aligned to the “off” bar.

The sum of synaptic inputs to each V1 cell was determined as

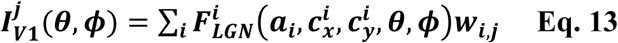

where *w_i,j_* is the strength of the synapse between LGN cell *i* and V1 cell *j*. V1 cell firing rates were modulated by lateral inhibition similar to that described for place cells in model two. The firing rate of a V1 cell was calculated as

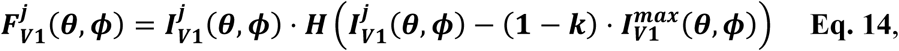

where 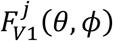 is the V1 cell’s firing rate at particular grating orientation and phase, *H* is the Heaviside function, and 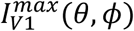 is the sum of excitatory input received by the most strongly excited V1 cell at orientation *θ* and phase *ϕ*. The parameter *k* is the fraction of 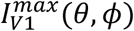 to which a V1 cell must be excited in order to fire. We set *k* = 0.10, as estimated for V1 (de Almeida, Idiart, & Lisman 2009b).

#### iv. Synaptic strengths

The strengths of LGN-cell-to-V1-cell synapses were sampled from the distribution described above for grid-cell-to-place cell synapses in model one. Each simulated recording session consisted of monitoring V1 cell activity as the grating stimulus rotated through 180-degrees. Following each full rotation, synaptic strengths at realized synapses between LGN and V1 cells were updated as 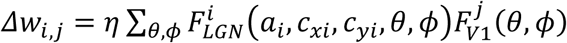 where the learning rate, *η*, equals one. After each strength update, we scaled the total strength of synapses converging onto each V1 cell by dividing by the sum of synaptic strengths and multiplying by the expected value of the sum of 130 random draws from the distribution of synaptic strengths.

#### v. Synapse turnover

Synapse turnover occurred following each full rotation of the stimulus grating. On average, 10% of active LGN cell inputs per V1 cell were replaced during each turnover event. This turnover rate was chosen because Tropea et al. (2010) (mouse, layer 5) and Yu, Majewska, & Sur (2011) (ferret, layer 2/3) observed that approximately 10% of V1 spines were replaced over a two-day interval. In our model, all realized synapses had an equal probability of removal. In contrast, the probability of a synapse forming between a particular LGN cell and a V1 cell was modulated by a Gaussian function (Eq. 15) so that synapses were more likely to form between neuron pairs with nearby receptive field centroids.

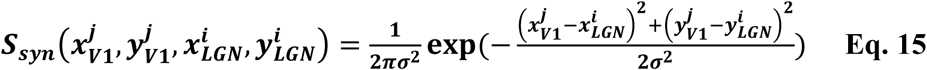

In equation 15, 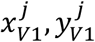 and 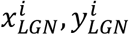 are the receptive field centroids of V1 cell *j* and LGN cell *i* respectively, and *σ* = 10. The probability of a synapse forming between a particular pre- and post-synaptic neuron pair was determined as

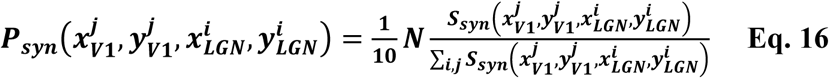

where *N* is the number of realized synapses onto the post-synaptic neuron. Thus, the average number of new synapses would be equivalent to the number of synapses removed per turnover event. Newly formed synapses took on random synaptic strength values sampled from the distribution described above.

#### vi. Receptive field analysis

“On” and “Off” stimulus patterns were generated as images with a single pixel set to one (“On”) or zero (“Off”) and all other pixels set to zero (“On”) or one (“Off”). One “On” and one “Off” stimulus was generated for each position in the 100x100 pixel visual field. For each V1 cell, separate “On” and “Off” receptive fields were calculated as a 2-dimensional map of synaptic input received by the cell when “On” and “Off” stimuli corresponding to each visual field position were presented to the network.

#### vii. Orientation preference analysis

V1 cell firing rates were averaged across grating phases for each grating angle. Each V1 cell’s preferred orientation was determined as the angle at which it exhibited the maximum average firing rate. The orientation preference shift on a given trial was calculated for all V1 cells as the difference angle between the cell’s preferred orientation on trial zero and a later trial.

## Results

### Nascent synapses are recruited into spatial memories

The crux of our hypothesis is that synapses that survive turnover provide a teaching signal for newly formed synapses. To illustrate this process, we first constructed a simple firing rate model (Fig. 1A) of a single place cell in CA1 driven by many grid cells in entorhinal cortex as a simulated animal runs in a linear track environment. In this model, the entorhinal cortex contains 10000 grid cells; however, only 1200 grid cells are synaptically connected to the place cell at any one time. We chose to use 1200 connections to approximate the total number of spatially modulated cortical inputs to a pyramidal cell in CA1 (see Materials and Methods). Each grid cell could provide at most a single synaptic connection to the place cell, and synaptic strengths were sampled from the empirically determined distribution (Eq. 4 & 5) (de Almeida, Idiart, & Lisman, 2009). In rodents, grid cells are active in evenly spaced grids across an environment, and the orientation and width of the grid vary by cell (de Almeida, Idiart, & Lisman, 2009). To capture these characteristics, we simulated grid cell firing rates as positive periodic functions of position (Fig. 1B) (Eq. 1) and assigned each grid cell a different grid rotation, phase, and period. When running a simulation, the linear environment was divided into 100 equally spaced discrete positions, and the place cell’s total synaptic input was calculated at each position (Eq. 2). To ensure that the place cell would always have a place field (Fig. 1C), we set the place cell’s threshold dynamically such that the cell would be active at the 10 positions where it received the greatest synaptic input. At these positions, the firing rate was equal to the sum of synaptic inputs.

**Figure 1.**
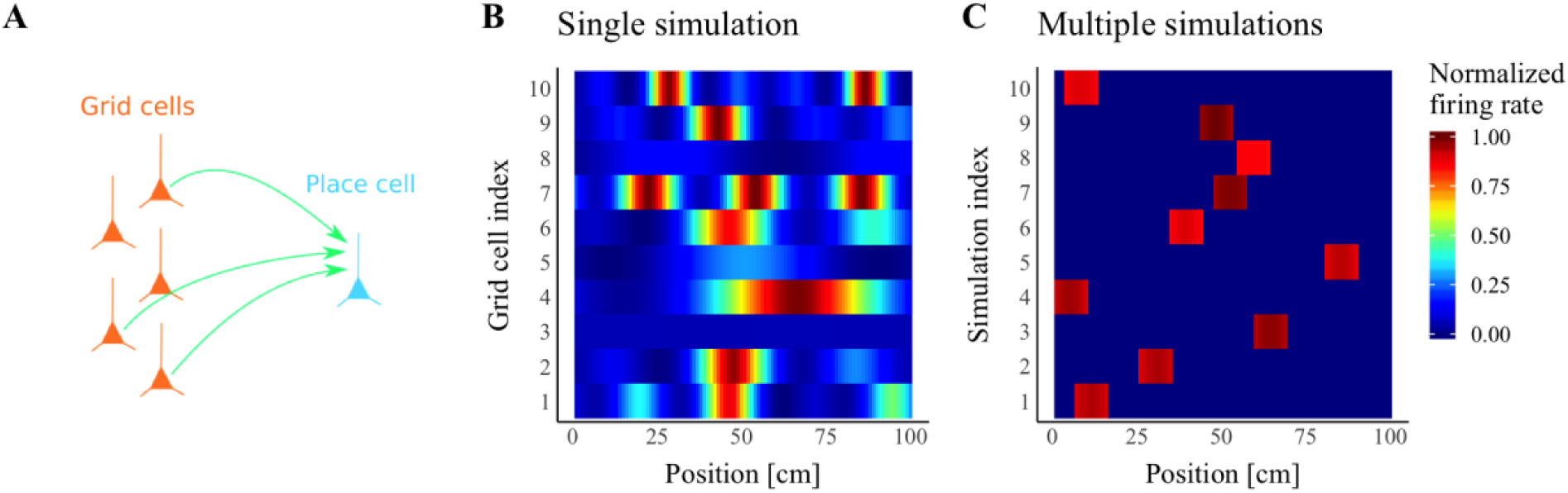
A feedforward convergent place-cell model of the grid-to-place-cell transformation. (**A**) Each individual place cell (blue) in CA1 receives synaptic input (green arrows) from many grid cells (orange) in entorhinal cortex. (**B**) As the simulated animal moves through a linear enclosure, the firing rates of the grid cells vary as periodic functions of position such that most grid cells have multiple peak firing locations (Each row depicts the activity of a single representative grid cell). (**C**) As the simulated animal moves, each place cell’s firing rate varies as the thresholded sum of its synaptic inputs from grid cells. The result of this process is that most place cells have a single firing field in the enclosure. (Each row depicts the activity of a single place cell receiving inputs as shown in A).

Each simulation included two sessions on the linear track, and sessions were separated by between one and 30 days. Each session was subdivided into an early phase and a late phase. The early phase was used to inform learning, and the late phase was used to measure place fields after initial learning but before synapse turnover. During both phases, the animal visited each position on the track with equal frequency. At the end of each early phase, we updated the strengths of synapses onto the place cell according to Hebb’s rule (Eq. 6) and implemented homeostatic scaling by divisive normalization (see Materials & Methods). Although LTP is not required for place field formation (Kentros et al., 1998; Agnihotri et al., 2004), we chose to have learning occur in the middle of each session because place cell firing can induce LTP within minutes (Isaac et al., 2009) and the spatial firing patterns of place cells in NMDAR1 knockout mice are more variable than those in wild type mice in novel environments (Cabral et al., 2014), suggesting that learning normally occurs early during exploration.

Synapse turnover occurred during the time between sessions. To simulate synapse turnover, we removed between 10% (approximately one day of turnover [Attardo, Fitzgerald, & Schnitzer, 2015]) and 100% (30 or more days of turnover [Attardo, Fitzgerald, & Schnitzer, 2015]) of the existing grid-cell-to-place-cell synapses, and we created an equal number of new synapses between the place cell and randomly selected, previously unconnected grid cells. The strengths of newly formed synapses were sampled from the same distribution that gave rise to the initial set of synaptic strengths.

To assess the effect of synapse turnover on the integrity of a place cell’s firing field, we compared that place cell’s response during the late phase of session one to its response during the late phase of session two. We defined integrity as the Pearson’s correlation between these responses. We performed 1000 separate simulations, varying the amount of synapse turnover, and found that integrity was high (with a median greater than 0.5) when as many as 70% of synapses were replaced during turnover (Fig. 2A). These data suggest that a place cell can maintain its firing field after the majority of its original synapses are turned over.

**Figure 2.**
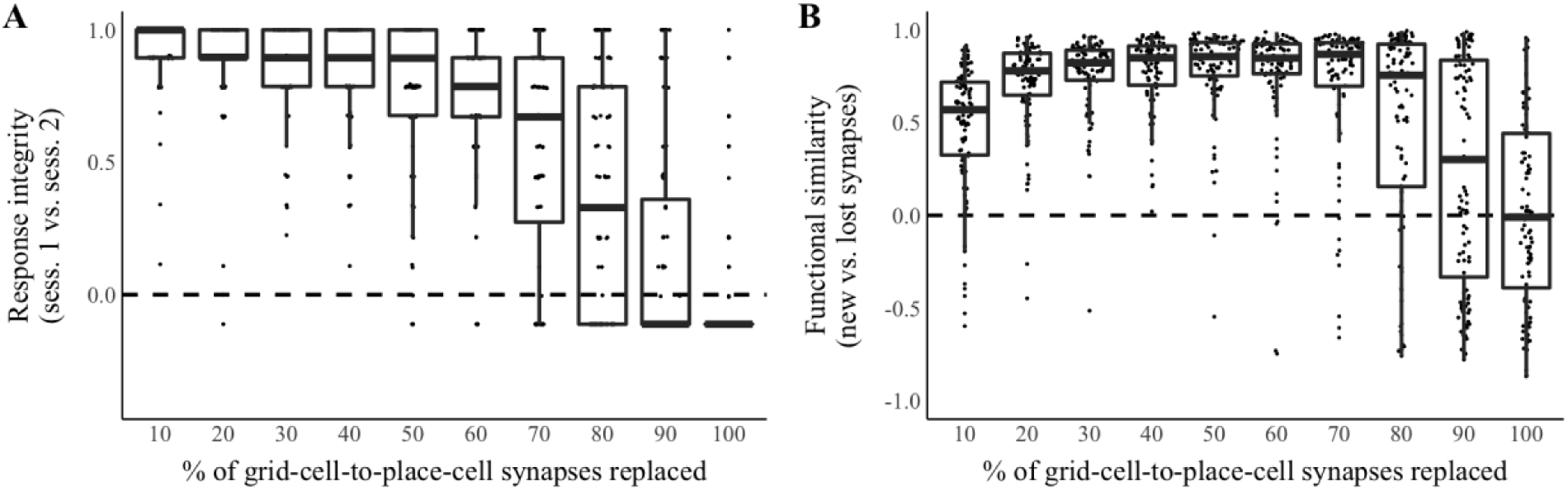
Retained synapses provide a teaching signal to nascent synapses. Activity in a single place cell model of the grid-cell-to-place-cell transformation was simulated over two sessions in a linear enclosure. Hebbian learning occurred during sessions, and 10% to 100% of synapses turned over between sessions. (**A**) Place cell response integrity score (Pearson’s correlation between the cell’s responses during the late phases of sessions one and two) versus the percent of synapses replaced between sessions. In these and all boxplots, hinges represent the first and third quantile, the thick bar represents the median, and the whiskers extend from the first quartile to the first quartile minus 1.5 times the interquartile range (IQR) and the third quartile plus 1.5 times the IQR. N=100 simulations per condition. (**B**) Synaptic functional similarity score (Pearson’s correlation between the sum of synaptic inputs by position during the late phase of session one at synapses removed due to turnover between sessions and the sum of synaptic inputs by position during the late phase of session two at synapses formed due to turnover between sessions) versus the percent of synapses replaced between sessions. N=100 simulations per condition.

Importantly, if retained synapses provide a teaching signal for new synapses, we expect that newly formed synapses will be functionally similar to synapses lost during turnover. To test functional similarity between new and lost synapses, we took the sum of synaptic input per position at 10% to 100% of grid-cell-to-place-cell synapses during the late phase of session one. We removed these synapses after session one and created an equal number of new synapses. We then took the sum of synaptic input per position at these new synapses during the late phase of session two. We determined a functional similarity score defined as the Pearson’s correlation between these sums of synaptic input at new and lost synapses. We found that functional similarity scores were high when up to 80% of synapses were replaced during turnover (Fig. 2B). These data suggest that a place cell’s latent position selectivity can provide a teaching signal for new synapses even after substantial synapse turnover.

In mice, blocking of the processes necessary for long-term potentiation when the animal investigates a novel environment disrupts the long-term stability of place fields associated with that environment (Kentros et al., 1998; Agnihotri et al., 2004). We therefore asked whether learning in the novel environment (session one of our paradigm) is required for the training of new synapses that is necessary for place-field stability in our model. To test this, we performed 200 additional simulations, alternately omitting or including the Hebbian learning step in session one. Omitting the Hebbian learning condition imitates the administration of a protein synthesis blocker or NMDA receptor antagonist in the novel environment. In these simulations, 10% of synapses were replaced between sessions one and two. Similar to *in vivo* reports (Kentros et al., 1998; Agnihotri et al., 2004), we observed a decline in place response integrity on session two when learning did not occur previously in the novel environment (*p* < .001, Wilcoxon rank sum test) (Fig. 3A) even though Hebbian learning could occur for the new synapses present in session two. In a further 200 simulations, we measured the functional similarity between new and lost synapses as described above while alternately omitting or including the Hebbian learning step in session one. We found that without learning during session one, functional similarity scores were not significantly different from zero (*p* = .887, Wilcoxon rank sum test) (Fig. 3B). Taken together, these data show that our model reproduces the experimental finding that an initial period of learning in a novel environment is required for the later recruitment and training of nascent synapses, leading to a robust long-term place memory.

**Figure 3.**
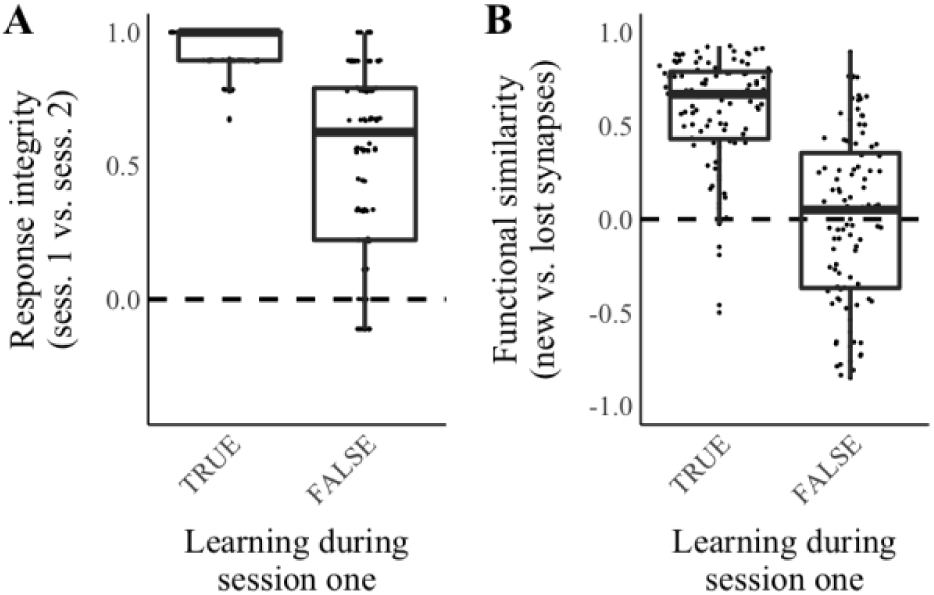
Training nascent synapses requires prior learning in the novel environment. Activity in a single place cell model of the grid-cell-to-place-cell transformation was simulated over two sessions in a linear enclosure. Hebbian learning occurred during sessions, and 10% of synapses turned over between sessions. In 50% of the simulations, Hebbian learning was omitted during session one, while still present for all synapses in session two. (**A**) Place cell response integrity score (Pearson’s correlation between the cell’s responses during the late phases of sessions one and two) versus a Boolean value indicating whether or not learning occurred during session one of each simulation. N=100 simulations per condition. (**B**) Synaptic functional similarity score (Pearson’s correlation between the sum of synaptic inputs by position during the late phase of session one at synapses removed due to turnover between sessions and the sum of synaptic inputs by position during the late phase of session two at synapses formed due to turnover between sessions) versus a Boolean value indicating whether or not learning occurred during session one of each simulation. N=100 simulations per condition.

As Hebbian learning is critical to the training of nascent synapses in our model, we decided to perform a parameter check to determine the range of learning rates over which such training is possible. To do this, we performed additional, two-session simulations as described above while varying the learning rate (*η* in Eq. 6) over a 10-million-fold range. Ten percent of synapses were replaced between sessions. We evaluated the functional similarity between new and lost synapses by determining a functional similarity score as previously described. We found that functional similarity scores increased rapidly as the learning rate increased from 1e-5 to 1e-3 (Fig. 4A). We chose to use a learning rate of 1e-4 for simulations described in this manuscript because 1e-4 was the smallest tested value that resulted in a functional similarity score significantly different from zero (*p* < .001, Wilcoxon rank sum test with Bonferroni corrections for multiple comparisons). In Figure 4B, we show the average absolute change in synaptic strength from the early to the late phase of session one at a variety of learning rates in the presence or absence of synaptic scaling. At learning rates below 1e-5, changes in synaptic strength are dominated by scaling. At a learning rate of 1e-4, the average change in synaptic strength is approximately 50% (41% with scaling; 62% without scaling) of the mean synaptic strength prior to learning (Fig. 4B, dashed line). At learning rates greater than 1e-3, with scaling, the average change in strength saturates at approximately 107% of the mean synaptic strength prior to learning. Changes in synaptic strength of 50% to 100% are consistent with biological data indicating that 10 stimulations of the pre-synaptic terminal can trigger a 100% increase in the average strength of CA3-CA1 synapses (Murthy, 1998). Further, we find that learning over two running sessions at a rate of 1e-4 preserves key properties of the synaptic strength distribution: unimodality and heavy-tailedness (Fig. 4C).

**Figure 4.**
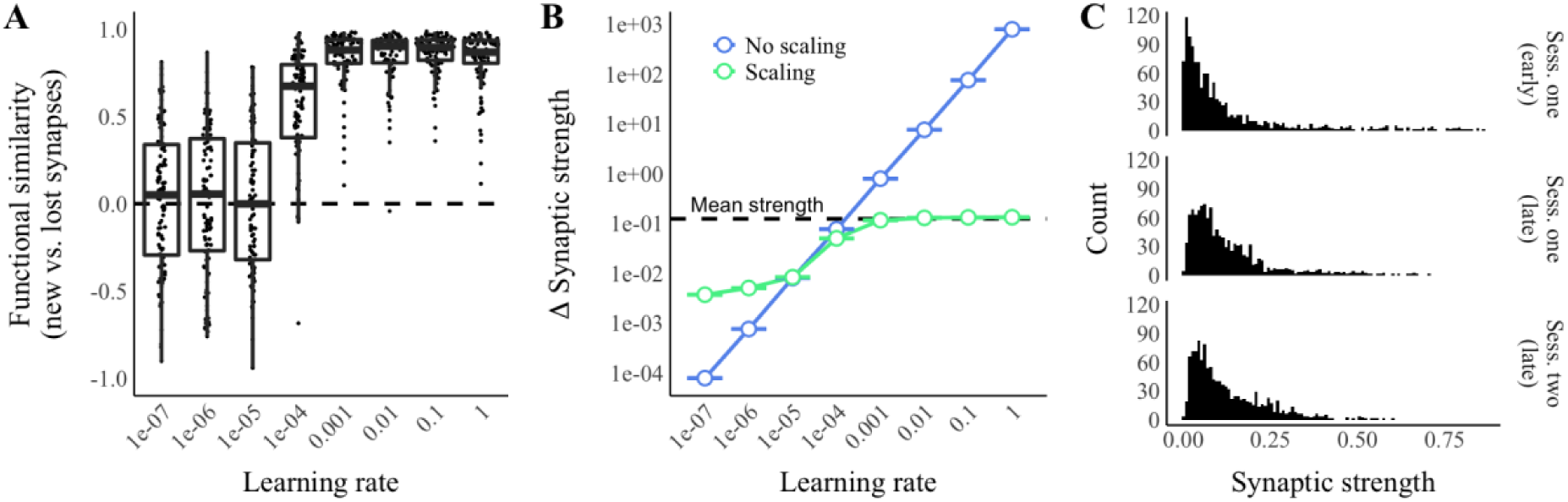
Effects of learning rate on the training of nascent synapses. Activity in a single place cell model of the grid-cell-to-place-cell transformation was simulated over two sessions in a linear enclosure. Hebbian learning occurred during sessions, and 10% of synapses turned over between sessions. (**A**) Synaptic functional similarity score (Pearson’s correlation between the sum of synaptic inputs by position during the late phase of session one at synapses removed due to turnover between sessions and the sum of synaptic inputs by position during the late phase of session two at synapses formed due to turnover between sessions) versus the learning rate. N=100 simulations per condition. (**B**) Absolute changes in synaptic from the early to the late phase of session one versus learning rate in the presence or absence of scaling. Error bars represent the mean +/- SEM across the strengths of synapses pooled from 10 simulations. The dashed line represents the overall mean synaptic strength prior to learning. (**C**) Representative distributions of synaptic strengths across all synapses by experimental stage in one simulation with a learning rate of 1e-4.

### Hebbian learning stabilizes place fields

It is important to evaluate place code stability in the context of the larger CA1 network because interneuron-mediated feed-back inhibition may control which cells fire in a particular environment and location (de Almeida, Idiart, & Lisman, 2009). To simulate the CA1 network, we turned to a previously established model defined by de Almeida, Idiart, and Lisman (2009). This model is identical to the single place cell model we described in the previous section, except for the addition of feed-back inhibition and an expanded pyramidal cell population (2000 cells) in CA1 (Fig. 5). This population model makes three assumptions about feed-back inhibition: 1) that pyramidal cells form convergent, excitatory synapses onto interneurons (Paz & Huguenard, 2015; English et al., 2017), which in turn form divergent, inhibitory synapses onto pyramidal cells (Bezaire & Soltesz, 2013); 2) that interneurons fire in response to a single excitatory synaptic potential (de Almeida, Idiart, & Lisman, 2009); and 3) that interneuron-to-place-cell synapses are strong enough to consistently suppress pyramidal cell firing (de Almeida, Idiart, & Lisman, 2009). These assumptions imply that, for a given cycle of the gamma oscillation, the first CA1 pyramidal neuron to fire will cause an interneuron to fire, which will in turn suppress the activity of any pyramidal neurons that do not fire before receiving synaptic input from the interneuron (de Almeida, Idiart, & Lisman, 2009). Since the neurons receiving the greater excitatory input will overcome the initially decaying inhibition and fire earlier in a gamma cycle, we can approximate this interneuron/pyramidal neuron interaction with a rule stating that CA1 pyramidal cells are only able to fire if they are excited to within some fraction (***k*** in Eq. 3) of the excitation received by the most excited cell (de Almeida, Idiart, & Lisman, 2009).

**Figure 5.**
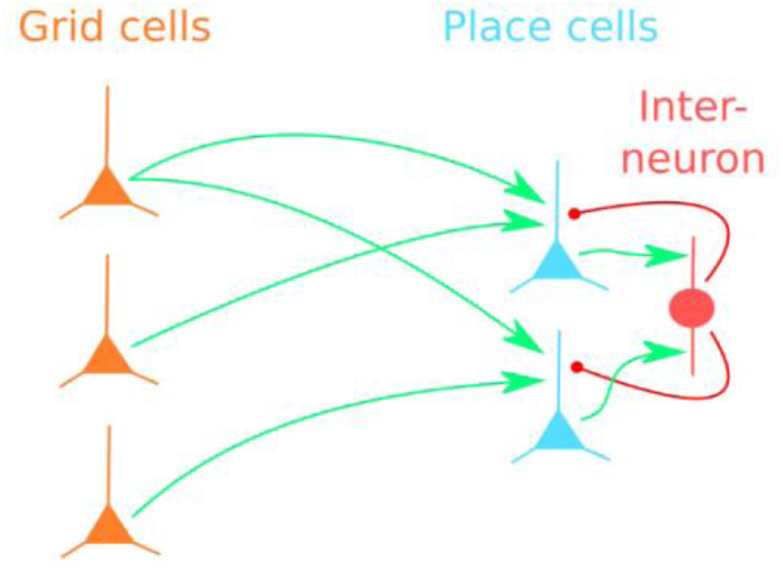
A multi-place-cell, feed-back-inhibition model of the grid-cell-to-place-cell transformation. Many place cells (blue) in CA1 receive synaptic input (green arrows) from many grid cells (orange) in entorhinal cortex. Place cells in CA1 form convergent, excitatory connections (green arrows) onto interneurons in CA1, which in turn form divergent, inhibitory connections (red arrows) onto the place cells. Interneuron-mediated feed-back inhibition controls which cells fire at particular locations in an enclosure.

Each of our simulations included 61 running sessions in the linear enclosure representing 61 days. The first running session is analogous to an experience in a novel environment. Subsequent sessions can be considered either as repeated experiences in the same environment or as offline events during which the original sequence of grid cell activity is replayed. As before, sessions were divided into an early phase and late phase with learning occurring at the transition between phases according to the Hebb’s rule (Eq. 6). To prevent synaptic strengths from increasing without bound, we simulated homeostatic synaptic scaling using divisive normalization (see Materials & Methods). Synapse turnover occurred at the end of each session. The rate of turnover was set at 9.5% (see Materials & Methods), the amount of synapse turnover expected during one day in mouse CA1 (Attardo, Fitzgerald, & Schnitzer, 2015). We defined place fields as continuous regions of the track, at least five centimeters in length, in which a cell’s average firing rate was within 80% of its maximum firing rate. During the late phase of each session, we extracted the place field centroids of all CA1 cells with place fields. We excluded cells with more than one place field.

To measure the effect of synapse turnover on place code stability, we compared the centroids of place fields found on the first session with the centroids of place fields found on later sessions. Comparisons were made within cell so that we were able to measure the drift in the position of an individual place cell’s place field. We defined drift as the absolute value of the within-cell difference between place field centroids on session one and a later session. If a cell did not have a place field on a particular session, that cell’s drift was undefined for that session. We ran 10 independent simulations and found that mean place field drift on the final session was significantly and dramatically reduced compared to 10 control simulations in which Hebbian learning steps were omitted (*p* < .001, Wilcoxon rank sum test) (Fig. 6A,B). These data suggest that Hebbian learning of synaptic strengths is sufficient to stabilize the place fields of individual place cells in a network, even after all original synapses are replaced.

**Figure 6.**
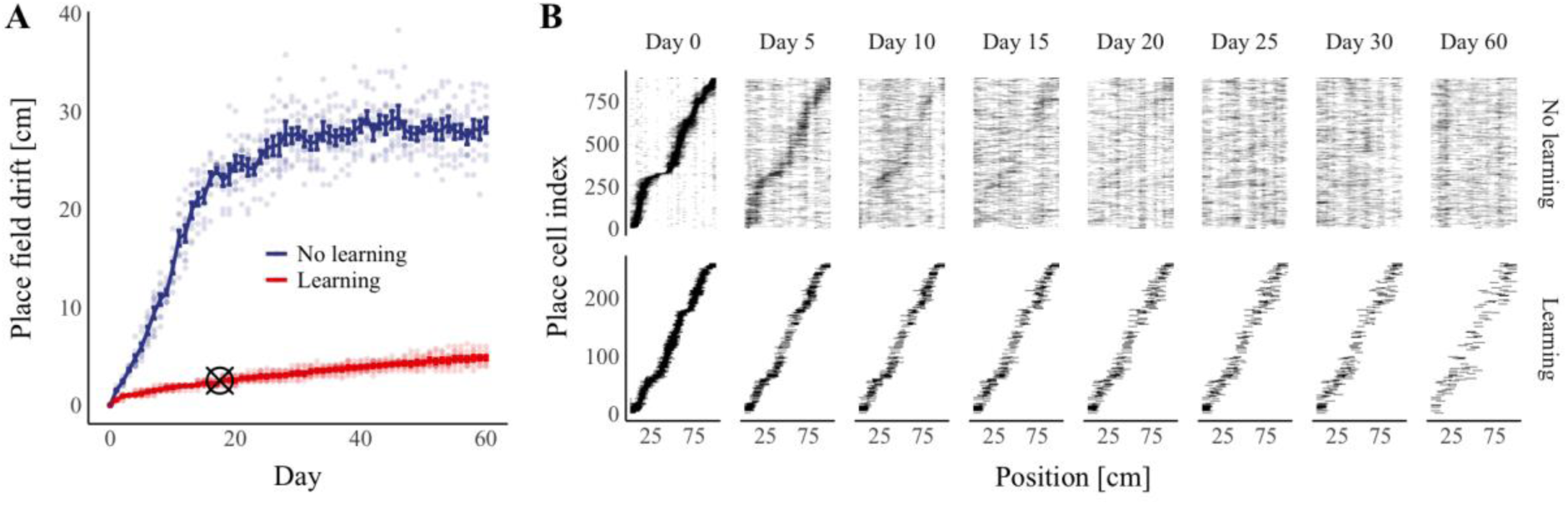
Hebbian learning improves place field stability. Activity in a feed-back-inhibition, grid-cell-to-place-cell model was simulated over a period of 61 days with both Hebbian learning and synapse turnover. (**A**) Median place field drift from day zero versus time and learning condition. Hebbian learning steps were omitted in the “No learning” condition. Error bars represent the mean +/- the standard error of the mean for median drift across simulations. Points represent median drift in individual simulations. Crosshairs indicate the experimentally determined median drift across days five through 30 in five day increments; crosshairs are positioned on the x-axis at the center of this time window (17.5 days). N=10 simulations per condition. (**B**) Representative place cell firing fields sorted by place field centroid on day zero. Black areas represent regions of the enclosure in which a cell was active.

As our choice of learning rate (1e-4) was motivated by a parameter check (Fig. 3A) rather than biological data, we decided to investigate the robustness of our model to changes in the learning rate. In our model, the overall distribution of synaptic strengths is relatively unchanged by learning (Fig. 3B). Therefore, the learning rate can be seen as modulating the degree to which a synapse’s strength after learning depends on its strength prior to learning. We performed an additional 80 simulations as described above, while varying the learning rate (*η* in Eq. 6) over a 10-million-fold range. We found that increasing the learning rate from 1e-6 to 1e-5 produced a dramatic reduction in the rate of place field drift (Fig. 7A), and all examined learning rates greater than 1e-5 resulted in less than 10 centimeters of average drift over 61 sessions (Fig. 7A). Our simulated results compare favorably to calcium imaging data showing 3.5 centimeters of median drift when aggregating data from days 5 to 30 in five-day increments (Ziv et al., 2013). Performing the same calculations over simulations performed at a learning rate of 1e-4, we observed 2.5 centimeters of median drift.

To further evaluate the effect of learning rate on place field properties, we measured the total number of place cells with place fields on each day (Fig. 7B), as well as the probability of place cell recurrence (Fig. 7C). We defined the probability of recurrence as the fraction of place cells with place fields during session one and a later session. We found that the total number of place cells and the number of recurring place cells followed a similar trend, decreasing as the learning rate increased from 1e-7 to 1e-5, then increasing towards an apparent asymptote as the learning rate increased from 1e-5 to one. Calcium imaging data indicate that approximately 20% of CA1 cells have a place field on a given session, while dividing the session into left and right moving trajectories reveals that approximately 10% of CA1 cells have a place field along a particular trajectory (Ziv et al., 2013). In our simulations, with a learning rate of 1e-4, an average of 7% of cells have place fields each day, an amount that is comparable to the experimentally determined percentage of cells with place fields along one trajectory.

To interpret the non-monotonic relation between the total number of place cells and the learning rate, we inspected the distribution of total synaptic input across CA1 cells in our model at each learning rate (Fig. 7D). Without learning, total input followed a Gaussian distribution. When learning was introduced, two clusters emerged in the distribution of total input. The first cluster included the majority of cells and followed the initial Gaussian distribution, while the second cluster exhibited greater total synaptic input. The emergence of the second cluster corresponds to an increase in the excitation of the most excited cell. Thus, fewer cells fired prior to the arrival of feedback inhibition, reducing the number of active place cells. With higher learning rates, the number of units in the second cluster increased. Thus, more cells received a similar amount of excitation to the most excited cell, and more cells were able to fire before the arrival of feedback inhibition.

**Figure 7.**
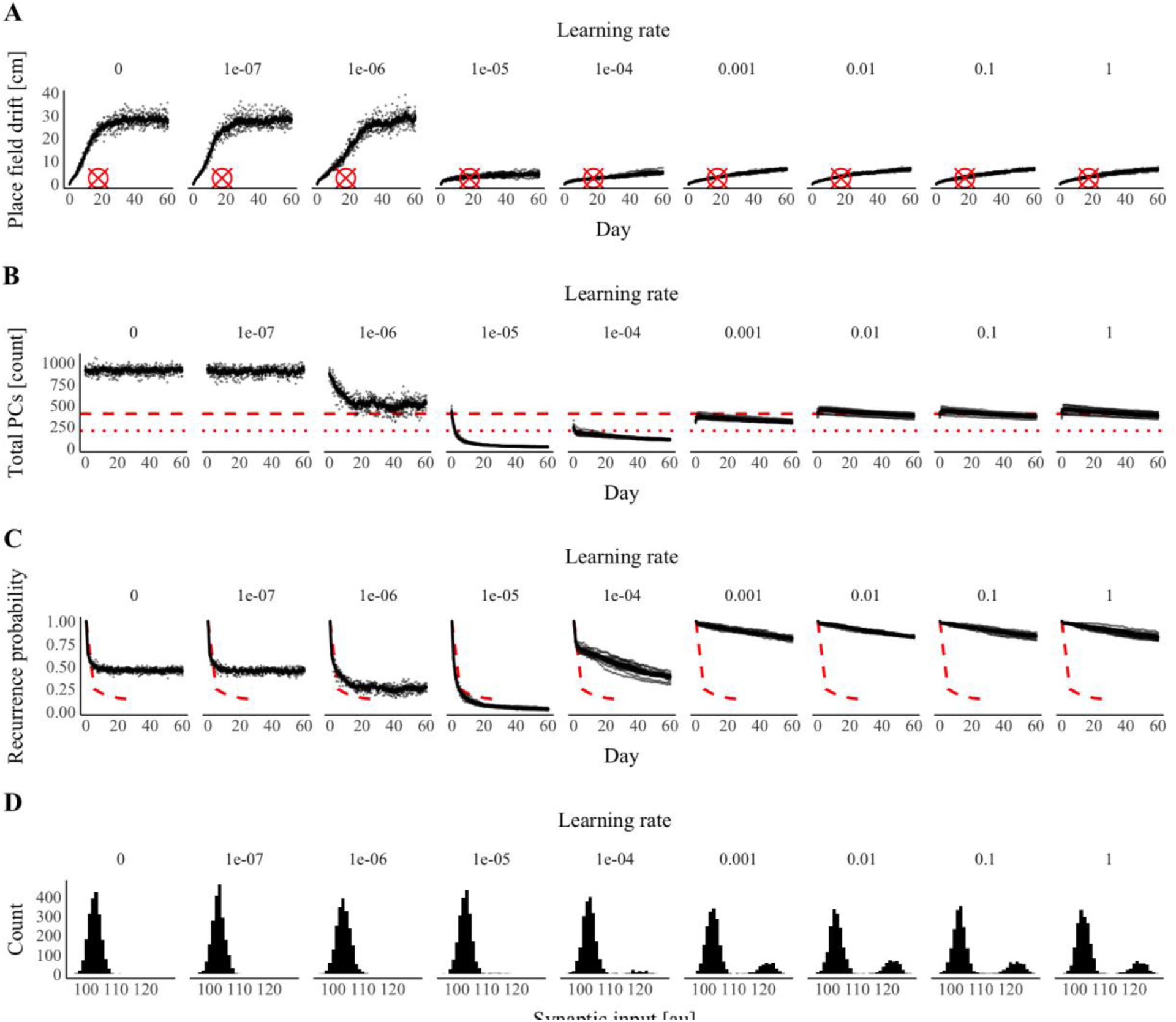
Effects of learning rate on place code properties. Activity in a feed-back-inhibition, grid-cell-to-place-cell model was simulated over a period of 61 days with both Hebbian learning and synapse turnover. The learning rate varied over a 10-million-fold range across simulations. (**A**) Median place field drift from day zero versus time and the learning rate taken during the late phase (after learning) on each day. Crosshairs indicate the experimentally determined median drift over days five through 30 in five day increments. Crosshairs are positioned on the x-axis at the center of this time window (17.5 days). (**B**) The total number of place cells (CA1 cells with place fields) versus time and the learning rate taken during the late phase (after learning) on each day. Dashed lines indicate the experimentally determined number of place cells per 2000 cells when the animal is moving in one direction (small dashes, 10%) or either direction (large dashes, 20%). (**C**) The probability of place cell recurrence (the fraction of CA1 cells with place fields on both day zero and a later date) versus the time and the learning rate taken during the late phase (after learning) on each day. Dashed lines indicate experimentally determined values from Ziv et al. (2013). (**A-C**) N=10 simulations per learning rate. (**A-C**) Data from the same set of simulations were used across panels. (**A-C**) Error bars represent the mean +/- the standard error of the mean across simulations, and points represent values from individual simulations. (**D**) Histograms of mean synaptic input per cell across positions on day 60 at several learning rates.

One caveat of our modeling approach thus far is that we did not account for the effects of irrelevant learning. For example, between running sessions, a mouse could be exposed to additional environments. Learning in these environments might interfere with previously stored memories. To address this complication, we asked if our model would be able to sustain place fields across multiple enclosures or if learning in a new environment would disrupt previously established place codes. To create a new linear enclosure, we shuffled the periodic firing rate functions associated with the grid cells. This procedure was motivated by the assumption that the firing fields of a grid cell in two environments are unrelated (de Almeida, Idiart, & Lisman, 2009). Our simulations in multiple enclosures were identical to our simulations in a single enclosure, except that three running sessions, in three separate environments, were experienced prior to each synapse turnover event. As before, learning occurred in the middle of each session. To assess place field stability across multiple environments, we quantified place field drift in each enclosure independently. We found that drift on the final session was significantly reduced in all three environments compared to control simulations with no learning (all *p* < .001, Wilcoxon rank sum tests with Bonferroni corrections for multiple comparisons) (Fig. 8). Aggregating all three environments, drift on the final session did not differ significantly from drift in experiments without interference (“Learning” in figure 6; *p* = .058, Wilcoxon rank sum test). Further, the median drift over days five through 30 in five day increments was 2.5 centimeters, similar to the biological value of 3.5 centimeters (Ziv et al., 2013). These results suggest that Hebbian learning is sufficient to preserve place codes despite both irrelevant learning and synapse turnover.

**Figure 8.**
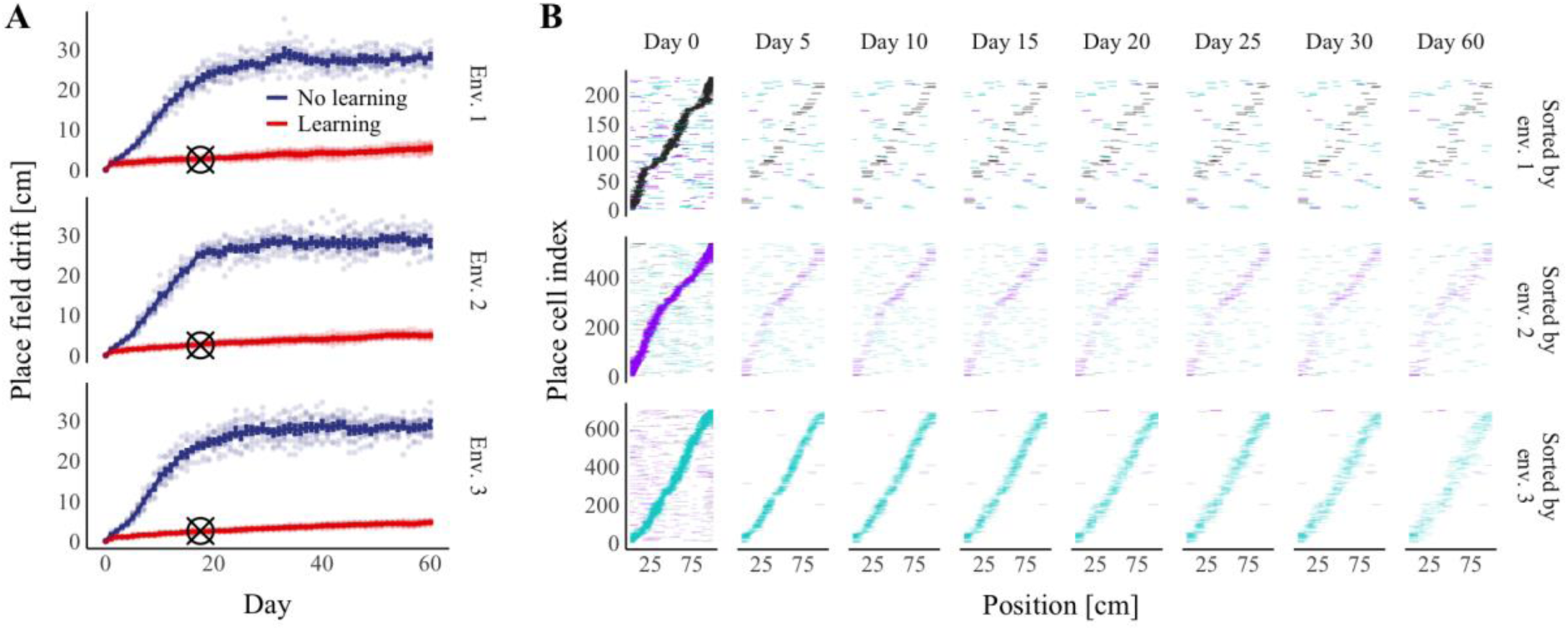
Hebbian learning improves place field stability across multiple environments. Activity in a feed-back-inhibition, grid-cell-to-place-cell model was simulated in three separate linear enclosure environments over a period of 61 days with both Hebbian learning and synapse turnover. (**A**) Median place field drift from day zero versus time in each environment. Hebbian learning steps were omitted in the “No learning” condition. Error bars represent the mean +/- the standard error of the mean for median drift across simulations. Points represent median drift in individual simulations. Crosshairs indicate the experimentally determined median drift across days five through 30 in five day increments; crosshairs are positioned on the x-axis at the center of this time window (17.5 days). N=10 simulations per condition. (**B**) Representative place cell firing fields from the “Learning” condition sorted by place field centroid on day zero in each of the three environments. Colored areas represent regions of the enclosure in which a cell was active. The color of a firing field represents the environment in which that field was observed (black, purple, and cyan correspond to environments one, two, and three respectively).

### Hebbian learning preserves memory with spontaneous reactivation

A condition of place field stability in the models we described thus far is that patterns of grid cell activity are replayed after initial learning. This condition is easily fulfilled if we assume that an animal will frequently and physically be re-introduced to an environment. Alternatively, patterns of grid cell activity may be replayed during rest, when the animal is not in the original environment. Several reports indicate that grid cell replay occurs during rest *in vivo* (Ólafsdóttir, Carpenter, & Barry, 2016; O’Neill et al., 2017; Ólafsdóttir, Bush, & Barry, 2018); however, it is not clear whether grid cell replay is intrinsic to cortex or requires hippocampal initiation (Ólafsdóttir, Carpenter, & Barry, 2016; O’Neill et al., 2017; Ólafsdóttir, Bush, & Barry, 2018). If grid cell replay is intrinsic to cortex, it may not be subject to degradation by synapse turnover, as cortical synapses appear to be much more stable than hippocampal synapses (Attardo, Fitzgerald, & Schnitzer, 2015). In contrast, if grid cell replay requires hippocampal initiation, it may depend upon the integrity of hippocampal replay (Ólafsdóttir, Bush, & Barry, 2018).

Hippocampal replay occurs during sharp wave ripples, highly synchronous events that occur during both active behavior and rest (Buzsáki & Mizuseki, 2015). Sharp wave ripples are thought to originate in CA3 (Buzsáki & Mizuseki, 2015). CA3 contains recurrent (feedback), excitatory connections between pyramidal cells and is often modeled as a Hopfield network (de Almeida, Idiart, & Lisman, 2007). Hopfield networks are recurrent, artificial neural networks that can reproduce each of many input patterns as a stable pattern of firing rates across neurons (fixed points) where the patterns are encoded in the strengths of connections between neurons (Cohen & Grossberg, 1983; Hopfield, 1983). If a Hopfield network is initialized in a random state, the network will converge to one of its stored patterns. Stored patterns in a Hopfield network can be maintained despite synaptic strength decay if the patterns are reactivated frequently and if Hebbian learning occurs during reactivations (Wittenberg, Sullivan, and Tsien 2002). Given the apparent importance of CA3 for memory reactivation *in vivo*, we asked how random synapse turnover would affect the stability of steady states in CA3-like Hopfield network models.

**Figure 9.**
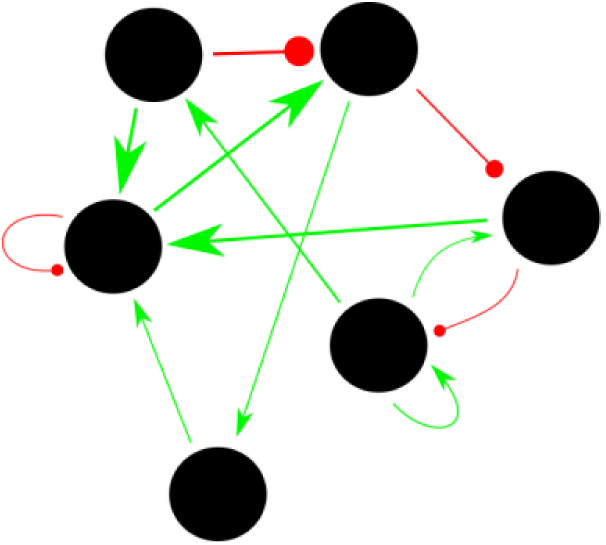
A sparse Hopfield memory network. Units (black) represent pyramidal cells in CA3. Units receive mono-synaptic input (arrows) from other units in the network. Synapses can be either excitatory (green) or inhibitory (red). Inhibitory synapses are taken to represent relays through fast-spiking interneurons. Synaptic strengths are initially set though a training procedure in which Hebbian learning occurs while the network receives strong external current injections that drive each unit toward a target firing rate. The trained state becomes an attractor of the system, and the network will converge to the trained state in the absence of external current sources.

Our Hopfield network models consisted of a variable number of units, sparsely connected by a variable number of monosynaptic synapses (Fig. 9). Firing rates were calculated as a sigmoidal function of synaptic input that could take on only positive values (Eq. 8). To train an individual pattern (see Materials & Methods), we clamped each cell’s firing rate at a target firing rate for the pattern and updated synaptic strengths according to a Hebbian learning rule that enabled both synaptic potentiation and depression (Eq. 9). Each simulation consisted of such an initial network training followed by 100 reactivation sessions with further learning, with the reactivation sessions preceded by synaptic turnover at a level corresponding to between one and 30 days of non-activation. Reactivation consisted of randomly initializing the membrane potentials of all units, allowing the network to converge to a steady state, and updating synaptic strengths according to the same Hebbian rule used for training (Eq. 9). During the synapse turnover between reactivation sessions, the connections between some units were broken, and an equal number of new connections were created.

**Table 1.** Parameters for Hopfield network experiments.

| Experiment | Figure/panel | Turnover rate | Probability of connection | # of units |
| --- | --- | --- | --- | --- |
| 1 | 10/A,D | [0%, 100%] | 0.2 | 100 |
| 2 | 10/B,E | 50% | (0.1, 0.8) | 100 |
| 3 | 10/C,F | 50% | 0.2 | Perfect squares in (16, 400) |
| 4 | 11 | [0%, 100%] | (0.1, 0.8) | Perfect squares in (16, 1024) |

We first performed simulations with a single stored pattern. We performed three experiments in which we alternately varied one of three parameters: the synapse turnover rate (Fig. 10A,D), the probability of connection between any two units (Fig. 10B,E), and the number of units in the network (Fig. 10C,F) while holding the remaining two parameters constant. When varied, parameters were randomly sampled in the ranges shown in Table 1, except for number of units which was randomly sampled from the set of perfect squares in the range. To assess pattern stability, we calculated an attractor integrity score as the Pearson’s correlation between the trained pattern and the network’s final state on the 100^th^ reactivation session. We found that the pattern integrity score decreased with synapse turnover (Fig. 10A) and increased with both the number of units (Fig. 10B) and the probability of connection (Fig. 10C).

**Figure 10.**
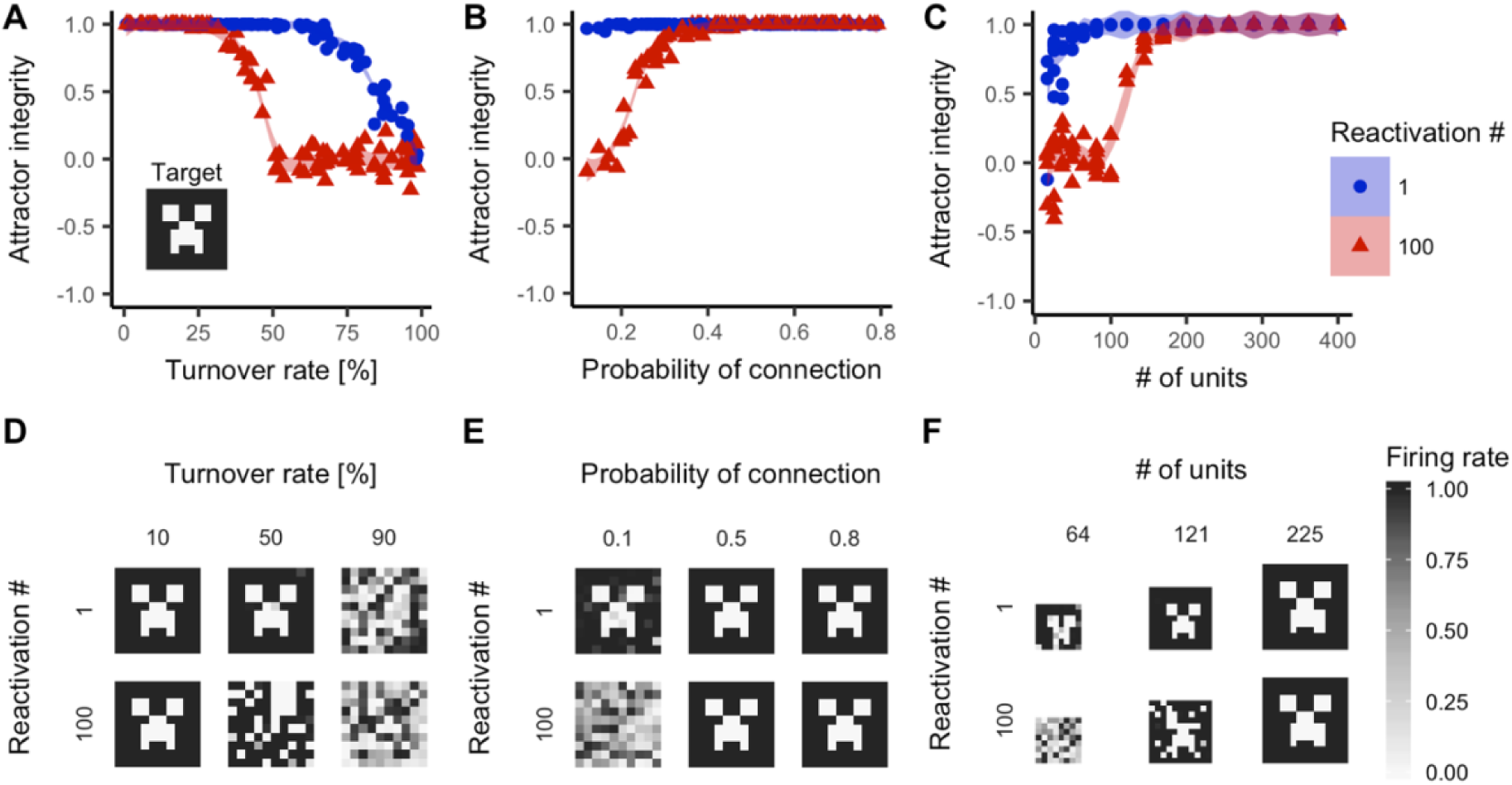
Network reactivation maintains attractor integrity in sparse Hopfield networks despite synapse turnover. Sparsely connected Hopfield networks were trained on an image (A: inset) and subjected to 100 sessions of reactivation and synapse turnover. An attractor integrity score (the Pearson’s correlation between the recalled attractor and the target firing rate vector) was calculated on the first and final reactivation. (**A**) Attractor integrity versus the percentage of synapses replaced per activation when the probability of connection and the number of units were constant at 100 and 0.2 respectively. (**B**) Attractor integrity versus the probability of connection when turnover rate and number of units were constant at 50% and 100 respectively. (**C**) Attractor integrity versus the number of units when the probability of connection and turnover rate were constant at 0.2 and 50% respectively. (**A,B,C**) Shaded areas represent 95% confidence intervals of locally weighted regression models. N=100 networks per panel. (**D,E,F**) Representative firing rates of networks as shown in A, B, and C.

To explore the extent to which the number of units and the number of connections could protect against synapse turnover, we performed 1000 additional simulations as described above while randomly varying all three parameters simultaneously over the ranges indicated in Table 1. For visualization, we used median in-degree (the number of inputs to a unit) as a stand-in for the probability of connection and the number of units, as in-degree is linearly dependent on both of these parameters. Integrity values of networks with identical in-degree and turnover rate were averaged, and integrity values of missing parameter combinations were estimated by bilinear interpolation (see Materials & Methods). We found that pattern integrity could be near one, indicating no degradation, in the presence of as much as 70% synapse turnover if the number of units and the probability of connection were sufficiently high (Fig. 11). These data demonstrate that reactivations and Hebbian learning can stabilize stored patterns in attractor networks such that the patterns can be recalled spontaneously despite synapse turnover.

**Figure 11.**
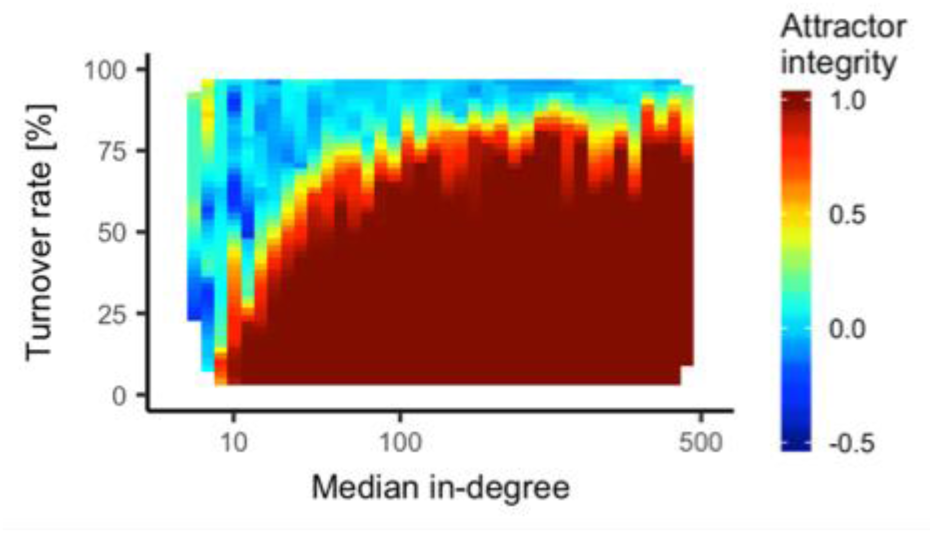
Attractors of large and more densely connected Hopfield networks are resistant to substantial synapse turnover. Sparsely connected Hopfield networks were trained on an image (Fig. 10A: inset) and subjected to 100 sessions of reactivation and synapse turnover. An attractor integrity score (the Pearson’s correlation between the recalled attractor and the target firing rate vector) was calculated on the final reactivation. The plot is a heat-map showing attractor integrity on reactivation 100 versus turnover rate and median in-degree when number of units, probability of connection, and turnover rate varied. N=1000 networks.

Next, we asked if multiple patterns could be preserved simultaneously in a Hopfield network undergoing synapse turnover. We initially turned to a method for multi-attractor stabilization described by Wittenberg, Sullivan, and Tsien (2002). In this method, multiple steady states are embedded in Hopfield network during training (see Materials & Methods). The model then experiences periodic reactivation sessions with synaptic strength decay between sessions. As demonstrated by Wittenberg, Sullivan, & Tsien (2002), when multiple attractors are present, it is necessary to reactivate each attractor in a directed manner, as random reactivation leads to a single attractor dominating network activity. In our simulations, during each reactivation session, the network is sequentially initialized in the vicinity of each attractor (see Materials & Methods). This is considered to be biologically reasonable because the activation of one pattern may bias a network towards a second pattern (Wittenberg, Sullivan, & Tsien, 2002). We performed simulations according to this protocol in networks with 400 units and a 0.5 probability of connection between any two units. We introduced synapse turnover between reactivation sessions and attempted to store three patterns simultaneously. We found that with 9.5% synapse turnover, as expected in a single day (Attardo, Fitzgerald, and Schnitzer, 2015), either all patterns were forgotten (integrity scores were low for all patterns) or a single pattern quickly dominated (integrity scores were high for one pattern and low for all others).

Cases where one pattern dominated suggested that patterns were not evenly represented after synapse turnover. However, features of pattern reactivation seen *in vivo* may act to rebalance stored patterns in the network. In resting rats, sharp wave ripples occur frequently (approximately 25 times per minute [Wiegand et al., 2016]). It may be that resting animals experience many replay events in short windows during which substantial synapse turnover is unlikely. We hypothesized that a process of reactivating each pattern many times would equalize the widths of the attractor basins, enabling multiple patterns to endure simultaneously. To test this, we performed additional simulations in which we reactivated each pattern sequentially by initializing the network in the vicinity of the corresponding attractor. We repeated this cycle of reactivations between one and 25 times per session, which is analogous to allocating at most one minute per day of resting time to each pattern. We also varied the learning time constant (*τ_w_*) in Eq. 9). Synapse turnover occurred only between sessions (not between sequences) at a rate of 9.5%. We assessed attractor integrity for each pattern after the final reactivation of that pattern on each session and found that frequency with which integrity was preserved (greater than .9) across all patterns on the final session tended to increase with the number of reactivation per session (Fig. 12). With 25 reactivations per session and *τ_w_* = 100, for example, all patterns were preserved over 100 sessions in 88% of networks (Fig. 12). These data suggest that bursts of reactivation can preserve multiple fixed points in a CA3-like network undergoing synapse turnover.

**Figure 12.**
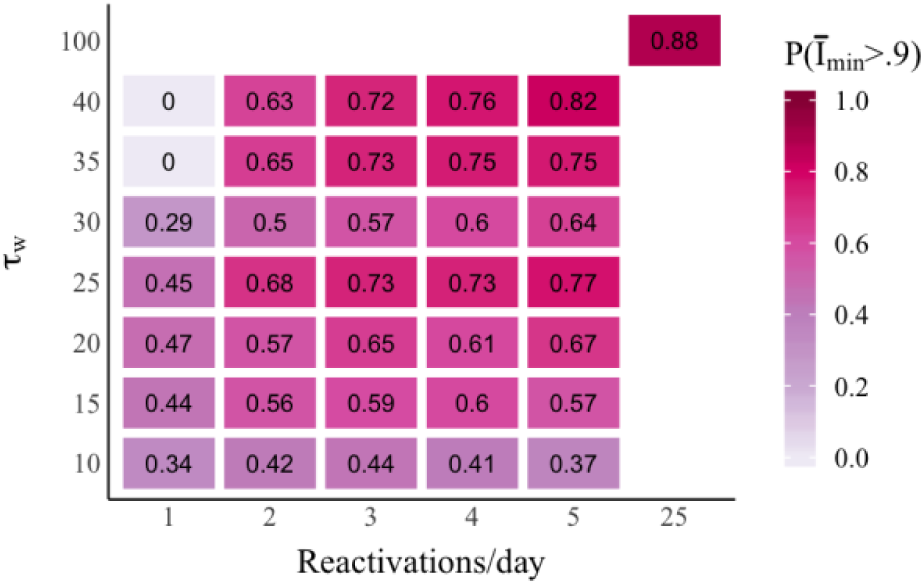
Bursts of sequential reactivation preserve multiple attractors in Hopfield networks undergoing synapse turnover. Sparsely connected Hopfield networks were trained on three randomly generated images and subjected to 100 days of reactivation and synapse turnover. During each day, a series of reactivations was performed by sequentially initiating the network in the vicinity of each stored pattern and incorporating Hebbian learning on the activity state reached with a learning time constant (*τ_w_*) of between 10 and 100. The series of reactivations was experienced between once or 25 times per day, and synapse turnover occurred between days. An attractor integrity score (the Pearson’s correlation between the recalled attractor and the target firing rate vector) was calculated for each pattern on the final reactivation of that pattern on each day. The plot shows the fraction of networks (out of 100 networks per parameter combination) for which the integrity score of the least well-preserved pattern was .9 or greater.

### Stable orientation tuning in an LGN-to-V1 model

So far, we have shown that Hebbian learning stabilizes memory in models of hippocampal function. However, our model may also help explain the stability of computation in non-hippocampal networks such as the mammalian visual system. For example, in V1, many cells firing selectively in response to visual stimuli at a particular orientation (Bonhoeffer & Grinvald, 1991). V1 cells with similar orientation preferences are clustered in pinwheel-patterned retinotropic maps (Bonhoeffer & Grinvald, 1991). Further, although synapses in V1 turn over in both developing and adult mammals (Holtmaat, Trachtenberg, & Wilbrecht, 2005; Stettler et al., 2006; Tropea et al., 2010; Yu, Majewska, & Sur, 2011), retinotropic maps can be stable throughout this period (Stevens et al., 2013).

To ask how activity-independent synapse turnover might affect the stability of orientation selectivity in V1, we constructed a simplified model of the LGN-center-surround-cell-to-V1-simple-cell transformation. In our model, visual stimuli were presented as a set of static sinusoidal gratings, across a square section of visual field, with gratings differing in their phase and angle of orientation. Visual input was received by 20000 LGN cells with on- or off-center difference-of-Gaussians receptive fields with random field centroids (see Materials & Methods). Each of 1000 V1 cells initially received excitatory synaptic input from 130 LGN cells. LGN inputs were initially arranged such that each V1 cell had a simple-cell-like receptive field consisting of an inhibitory central bar and two flanking excitatory bars (Fig. 13A, top) (Lee, Huang, & Fitzpatrick, 2016) (see Materials & Methods). The centroids and rotations of V1 cell receptive fields were assigned randomly. The firing rates of V1 cells were further modulated by lateral inhibition (see Materials & Methods).

We simulated network activity in sessions during which the static grating stimulus was presented at a set of angles sampled evenly over a 180-degree range. After each session, the strengths of LGN to V1 synapses were updated according to Hebb’s rule (see Materials & Methods), and an average of 10% of synapses were replaced (see Materials & Methods). Ten-percent is the approximate number of spines replaced over two days in adult mouse V1 (Tropea et al., 2010; Yu, Majewska, & Sur, 2011). In mice, spine turnover in V1 seems to be partially activity-dependent (Tropea et al., 2010); however, for simplicity, we assume that all rewiring is activity-independent. In addition, in mice, there appears to be a sizable population of stable spines in V1 (Holtmaat et al., 2005); however, we assume only a single population of unstable spines. In our model, the probability of removal was identical across all synapses, while the probability of synapse formation between two units was determined by a function that fell off with the distance between receptive field centroids. In total, we simulated 60 sessions, representing network activity over 120 days in two-day increments.

We found that the receptive fields and activity patterns of V1 cells were preserved between the first and final sessions. We visualized the receptive fields of individual V1 cells observing synaptic input at V1 in response to individual bright or dark spot stimuli at positions sampled evenly across the visual field (Fig. 13A). V1 receptive fields largely maintained their three-bar structure after 120 days. We also visualized the spatial tuning of synaptic inputs (Fig. 13B) and firing rates (Fig. 13C) for individual V1 cells across stimulus orientations at their preferred stimulus phase. We observed little change over time in the visual orientations producing peak synaptic input (Fig. 13B) and firing rates (Fig. 13C, D) for individual V1 cells. To quantify within-cell shift in orientation preference, we took the angle between the orientations of peak firing during the first and final session (Fig. 13E). In our model, mean shift was significantly reduced compared to a control in which final orientation preferences were shuffled (*p* < .001, Wilcoxon rank sum test). Meanwhile, the range of synaptic strengths remained similar across the simulated period (Fig. 13F). These data suggest that our model can help to explain stable neural representations in extra-hippocampal networks.

**Figure 13.**
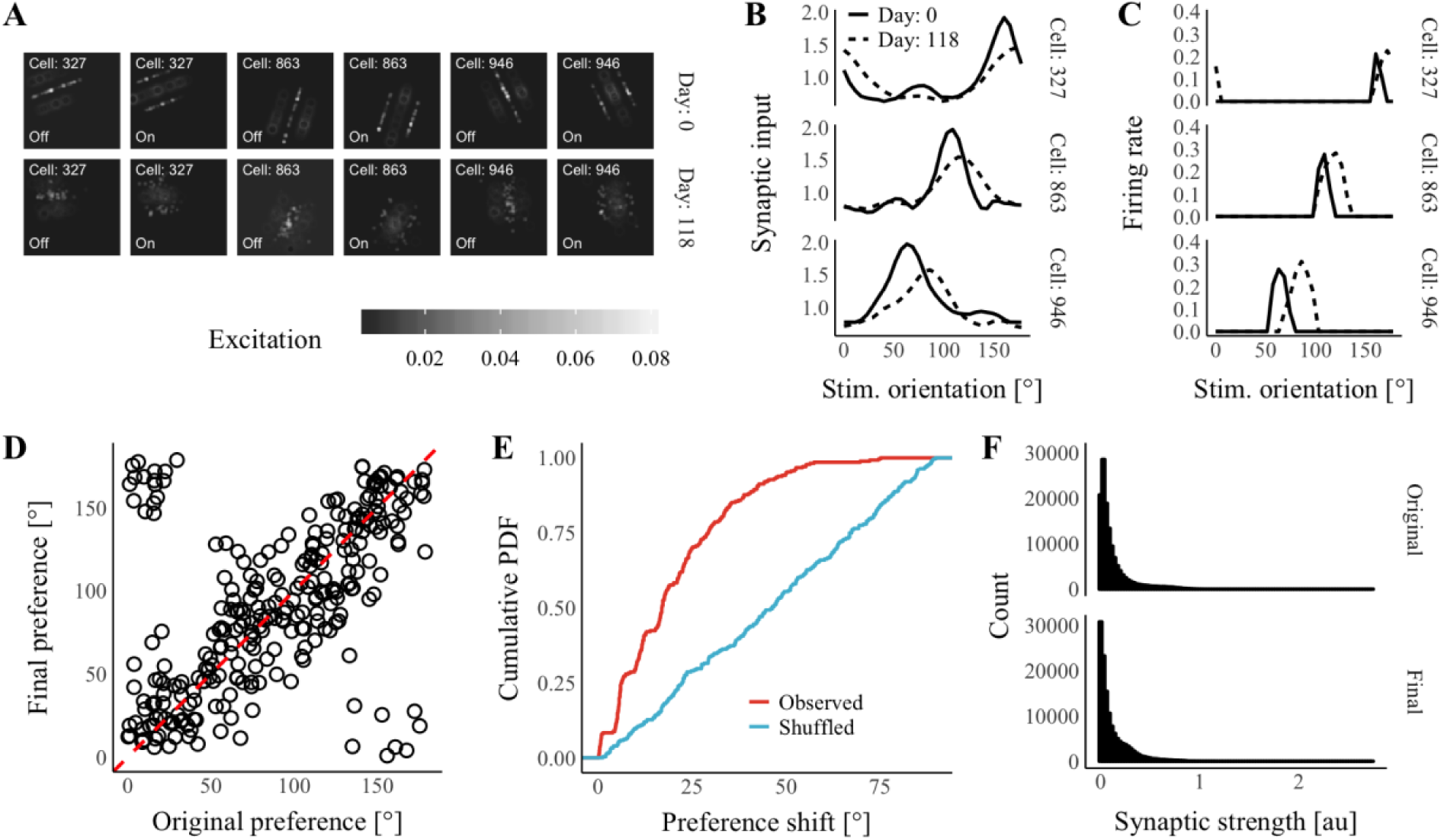
Persistent orientation tuning in an LGN-to-V1 model. (**A**) Visual receptive fields of three representative V1 cells on days zero and 118. Excitation is the amount of synaptic input received by the V1 cell in response to bright (“On”) or dark (“Off”) spot on the visual field. (**B**) Synaptic input received by cells in **A** at their most excited phase across orientations on days zero (solid lines) and 118 (dashed lines). (**C**) Firing rates of cells in **A** at their most-excited phase across orientations on days zero (solid lines) and 118 (dashed lines). (**D**) Original versus final preferred orientation for all V1 cells. (**E**) Cumulative probability density plot of observed orientation preference shifts (red) and a shuffled condition (blue). (**F**) Histograms of original and final synaptic strengths at realized synapses across all cells.

## Discussion

Recent research suggests that the neuronal connectomes of adult mammals are not as stable as once thought (DeBello & Zito, 2017). In regions such as the hippocampus, the majority of the synaptic architecture might be rewired in a period as short as one month (Attardo, Fitzgerald, & Schnitzer, 2015). Under these conditions, it is not clear whether or how memories and computational properties could be preserved in the long term. While complete synaptic turnover may take longer in cortical regions, the many decades of an animal’s lifetime are likely to prove to be too long for the set of connections formed in an early critical period to remain. Yet, for us to recognize a person we met or place we lived many decades earlier, a fraction of the original cortical representation should remain intact. Therefore, we sought to address the question of how representations could remain in the face of a long-term overhaul in the connectivity of a circuit, using a modeling approach. We found that Hebbian plasticity of synaptic strengths, in particular when combined with network reactivation, is sufficient to maintain network memory and function in the midst of random synapse turnover.

Specifically, in a network with many varying inputs, our model predicts that the synapses that survive turnover will provide a teaching signal to newly formed synapses. In time, all original synapses may turn over, but synapses that received the teaching signal will continue to pass that signal on as more new synapses are formed. Consistent with *in vivo* observations (Ziv et al., 2013), we found that replay and Hebbian plasticity led to stable place fields in models of hippocampal place cell networks. Further, we found that reactivations and Hebbian plasticity stabilized attractor states in CA3-like Hopfield networks. Together, these results suggest that random synapse turnover does not preclude long-term memory. To assess the generalizability of our model to non-hippocampal systems, we examined the stability of orientation tuning in a network model of V1 simple cells. In this model, we observed persistent orientation preferences despite activity-independent synapse turnover. This result suggests that, once established, simple learning rules may counteract effects of rewiring noise on orientation tuning maps.

Several pre-existing models suggest that memory and computation can outlive individual synapses (Poirazi & Mel, 2001; Hopfield & Brody, 2004; Knoblauch et al., 2014; Fauth, Wörgötter, & Tetzlaff, 2015; Eppler et al., 2015; Gallinaro & Rotter, 2017). However, our model is unique in two ways. First, in our model synapse stability and the location of new synapses are random. This is in contrast to models created by Poirazi and Mel (2001), Hopfield and Brody (2004), Knoblauch et al. (2014), and Fauth, Wörgötter, and Tetzlaff (2015), all of which involve activity-dependent synapse formation and/or elimination. Because we leave the control of synapse turnover to chance, our model is generalizable to neural structures such as the adult hippocampus in which mechanisms for activity-dependent wiring are poorly characterized. Second, our model requires only monosynaptic connections. This is in contrast to the model by Fauth, Wörgötter, and Tetzlaff (2015) in which information is represented by the number of realized synapses connecting neuron pairs. Because our model does not rely on compound connections, it can be extended to represent a variety of circuits with few potential connections per neuron pair.

An important open question is whether synapse turnover in the hippocampus is truly random. This is a difficult question to answer for several reasons. One reason is that some memory models require very few stable synapses, e.g. John Lisman favored a model in which only about 0.1% of spines must be stable (personal communication, 2017). Such a small stable population of dendritic spines would be below the threshold for detection by current live imaging technologies. Another reason is that spine stability might be related to spine size: LTP might lead to an expansion of the post-synaptic density and actin polymerization, resulting in an enlarged spine that is unlikely to collapse into the dendritic shaft (Bramham et al., 2010; Lüscher & Malenka, 2012). In agreement with this hypothesis, Pfeiffer and colleagues (2018), showed a small but significant relationship between spine lifetime and spine volume in adult mouse CA1. However, the high density of dendritic spines in the region, as compared to cortex, can lead to optical merging of nearby spines, increasing uncertainty around volume estimates (Attardo, Fitzgerald, & Schnitzer, 2015). To contribute to these discussions of hippocampal plasticity, we have shown that the place fields of individual place cells can be stable despite the ongoing, random replacement of grid cell inputs. Our results suggest that neither activity-dependent synapse stabilization nor an inherently stable synapse population is necessary to explain the observed persistence of hippocampal representations.

Our results generalize to any circuits of the brain undergoing synaptic turnover and Hebbian plasticity. Since some synapses have been observed to be stable, suggesting synaptic turnover is not an unavoidable biological necessity, one can question whether there is any advantage to circuits possessing connections that are not robust. Within cortical regions, which extract the important correlations from their inputs, such rewiring allows an animal to retrain when input statistics change. Within the hippocampus, the rewiring may allow for selective maintenance of memories, ensuring that only important or revisited experiences are connected to produce long-lasting episodic memories. In support of this idea, one theory of hippocampal and cortical memory is that hippocampus both learns and forgets quickly, while cortex learns more slowly but retains information longer (Lisman & Morris, 2001; Roxin & Fusi, 2013). According to this view, the memories in the hippocampus are slowly transferred to the cortex for long-term storage. These memories can then be cleared from the hippocampus, making way for new learning (Lisman & Morris, 2001; Lisman & Grace, 2005; Richards & Frankland, 2017). The regional timescales of memory retention in this model are correlated with the difference in apparent turnover dynamics between hippocampus and cortex: many cortical spines are long-lived compared to hippocampal spines. Our model predicts that hippocampal memories would eventually be cleared without reinforcement; however, learning during reactivation events could allow a memory to endure.

We focused on reactivation of neurons within a network possessing stable, persistent activity states, namely the Hopfield network, to explore the robustness of place fields in the absence of visitation of the corresponding environment. To date, the observed reactivations of neurons in CA3 have been in sharp wave ripples, which produce transient, sequential activity, as opposed to the persistent non-sequential activity of a Hopfield network. However, in both cases Hebbian plasticity has the similar effect of increasing the robustness of an attractor state. In our model, we found reactivation of attractor states leads to synaptic plasticity and is important for maintenance of place fields when a location is not visited frequently enough compared to the turnover rate. Likewise, sharp wave ripples can induce synaptic potentiation (Sadowski, Jones, & Mellor, 2016) and are important for spatial learning and memory (Jadhav et al., 2012).

In sum, we describe a model of memory preservation amidst synapse turnover that accounts for an apparent contradiction emerging from recent research on connectome stability and the persistence of memories encoded in neural ensembles. We show that memory and computation can be stabilized by Hebbian plasticity of synaptic strengths, even as the network is randomly rewired. In the future, it will be interesting to use the methods presented here to ask whether newly generated neurons are likely to be assimilated into preexisting engrams, or if they are solely useful in creating new memories.

